# *Bcl-xL* as a poor prognostic biomarker and predictor of response to adjuvant chemotherapy specifically in *BRAF*-mutant stage II and III colon cancer

**DOI:** 10.1101/207639

**Authors:** Philip D. Dunne, Helen G. Coleman, Peter Bankhead, Matthew Alderdice, Ronan T. Gray, Stephen McQuaid, Victoria Bingham, Maurice B. Loughrey, Jacqueline A. James, Amy M. B. McCorry, Alan Gilmore, Caitriona Holohan, Dirk Klingbiel, Sabine Tejpar, Patrick G. Johnston, Darragh G. McArt, Federica Di Nicolantonio, Daniel B. Longley, Mark Lawler

## Abstract

**Background:** BRAF mutation occurs in 8-15% of colon cancers (CC), and is associated with poor prognosis in metastatic disease. Compared to wild-type *BRAF* (*BRAFWT*) disease, stage II/III CC patients with *BRAF* mutant (*BRAFMT*) tumors have shorter overall survival after relapse; however, time-to-relapse is not significantly different. The aim of this investigation was to identify, and validate, novel predictors of relapse of stage II/III BRAFMT CC.

**Patients and methods:** We used gene expression data from a cohort of 460 patients (GSE39582) to perform a supervised classification analysis based on risk-of-relapse within *BRAFMT* stage II/III CC, to identify transcriptomic biomarkers associated with prognosis within this genotype. These findings were validated using immunohistochemistry in an independent population-based cohort of Stage II/III CC (n=691), applying Cox proportional hazards analysis to determine associations with survival.

**Results:** High gene expression levels of Bcl-xL, a key regulator of apoptosis, were associated with increased risk of relapse, specifically in *BRAFMT* tumors (HR=8.3, 95% CI 1.7-41.7), but not *KRASMT/BRAFWT* or *KRASWT/BRAFWT* tumors. High *Bcl-xL* protein expression in *BRAFMT*, untreated, stage II/III CC was confirmed to be associated with an increased risk of death in an independent cohort (HR=12.13, 95% CI 2.49-59.13). Additionally, *BRAFMT* tumors with high levels of *Bcl-xL* protein expression appeared to benefit from adjuvant chemotherapy (P for interaction =0.006), indicating the potential predictive value of Bcl-xL expression in this setting.

**Conclusions:** These findings provide evidence that *Bcl-xL* gene and/or protein expression identifies a poor prognostic subgroup of *BRAFMT* stage II/III CC patients, who may benefit from adjuvant chemotherapy.

**Key Message:**

- Using a combination of computational biology discovery and immunohistochemistry validation in independent patient cohorts, we show that high expression of the apoptosis regulator Bcl-xL is associated with disease relapse specifically within *BRAF* mutant stage II/III colon cancer.
- This data could enable tailored disease management to reduce relapse rates in the most aggressive subtype.

## Introduction

Results from a phase III trial (MRC COIN trial, n=1630) in metastatic CRC revealed that patients with *BRAF* mutant (*BRAFMT)* tumors have a significantly worse prognosis compared to patients with *KRAS* mutant (*KRASMT*) tumors or tumors with no detectable mutations in *KRAS* or *BRAF* (*WT/WT*).[1] Analysis of tumors in a cohort of 688 stage II and III colon cancer (CC) clinical trial samples (PETACC-3)[2] confirmed that *BRAF* mutation could identify a subgroups of patients with poorer OS <u>*after*</u> relapse, (i.e. when the patient had progressed to stage IV metastatic disease). Importantly however, *BRAF* mutation did not identify stage II/III patients with a higher rate of disease relapse, indicating the poor prognosis of *BRAFMT* disease was only evident in stage IV disease.

There is currently a lack of understanding of the biology that drives disease relapse, specifically within stage II/III *BRAFMT* disease. Therefore, we aimed to identify novel predictors of relapse for stage II/III *BRAFMT* CC, employing transcriptomic datasets for *in silico* discovery/initial corroboration, followed by subsequent validation of promising lead candidate(s) from bioinformatics analyses by immunohistochemistry analysis within a large population-based stage II/III *BRAFMT* CC study.

### Patients and methods

#### Transcriptional datasets

Gene expression profiles were downloaded from NCBI Gene Expression Omnibus (GEO) (http://www.ncbi.nlm.nih.gov/geo/) under accession number GSE39582 and GSE35602. As detailed in Supplementary Figure 1, GSE39582 [3, 4] contained 460 stage II/III CC profiles which had relapse data available. For initial biomarker discovery, the “Prognostic Subset” contained untreated stage II/III patients stratified into high-risk (if the patient relapsed within 36 months) or low-risk (if there was no relapse). The “Initial Consolidation” contained all stage II/III patients with relapse information and mutational data (n=417), which included *BRAFMT* (n=41; 24 of which will have been used already in the prognostic subset), *KRASMT* (n=166) or *WT/WT* (n=210) (Supplementary Figure 1).

**Figure 1:**
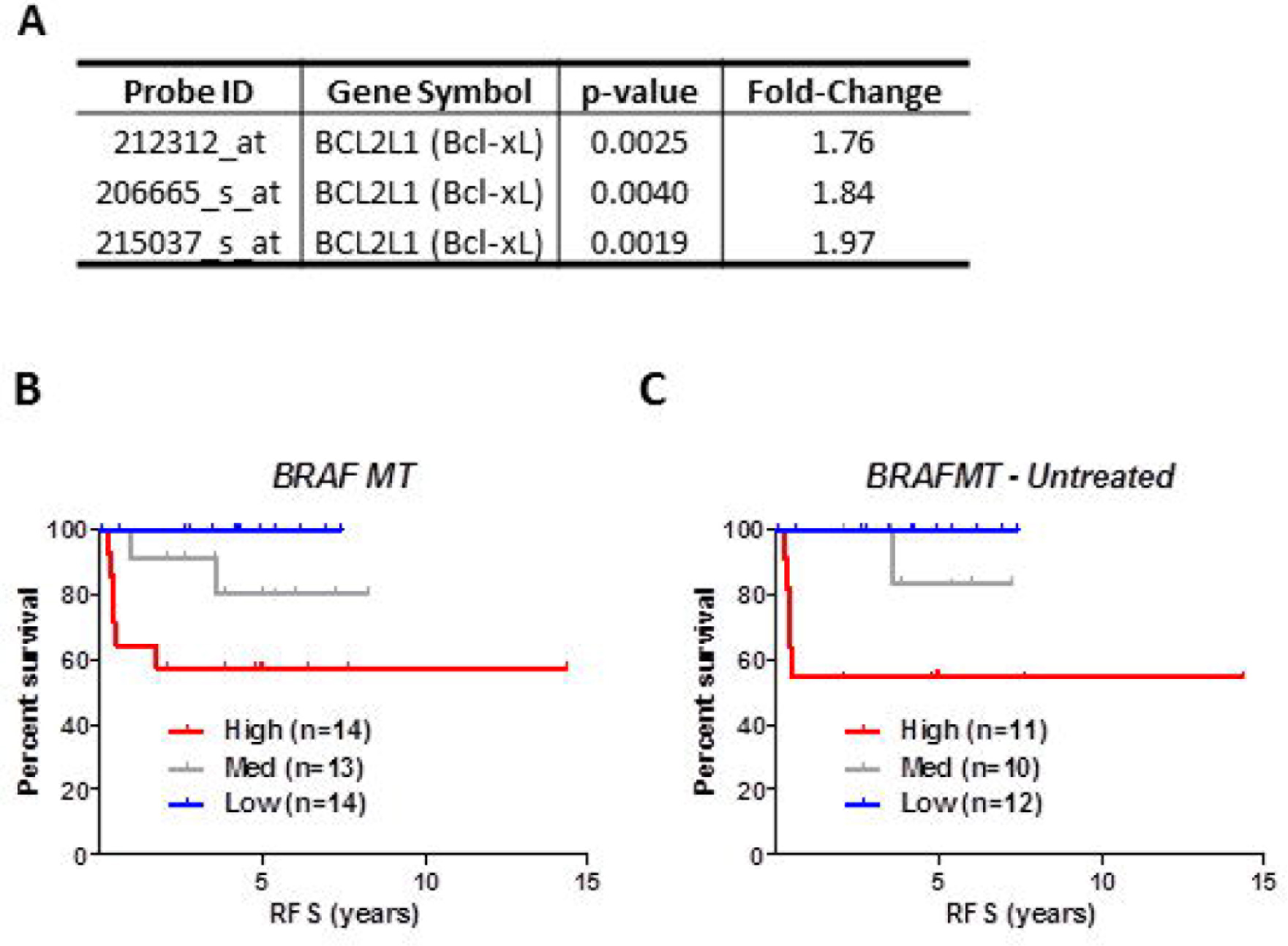
Relapse risk analysis of *BRAFMT* tumors indicates that *Bcl-xL* gene expression is associated with prognosis in *BRAFMT* tumors. **A.** BCL2L1 (*Bcl-xL*) was represented by 3 individual probesets in relapse risk analysis in *BRAFMT* tumors. **B + C.** Relapse-free survival (RFS) curve using Kaplan-Meier estimation in the “Initial Consolidation” dataset comparing tertile stratified *Bcl-xL* gene expression levels in all *BRAFMT* (A) and untreated *BRAFMT* (B) stage II/III CRC patients. Unadjusted and adjusted HR statistics are detailed in Table 3.

#### Transcriptional analysis

Partek Genomics Suite was employed for dataset analysis. Differentially expressed probesets which had a fold-change +/− 1.75 fold and p-value <0.005 were defined using analysis of variance (ANOVA) of supervised risk groupings in both the *BRAFMT* and *KRASMT* subgroups separately. Gene Set Enrichment Analysis was accessed (GSEA; http://software.broadinstitute.org/gsea/index.jsp) and the Microenvironment Cell Populations-counter (MCP) was accessed via the https://doi.org/10.5281/zenodo.61372 link.

### Bcl-xL Immunohistochemistry (Bcl-xL IHC)

*Bcl-xL* IHC antibody (Cell Signaling Technology, MA, United States) (*Bcl-xL* (54H6) Rabbit mAb #2764) was employed at 1:250 dilution with epitope retrieval solution 2 pretreatment for 30 minutes, visualized with diaminobenzidine, counterstained with hematoxylin, and mounted in DPX.

#### Independent stage II/III CC Northern Ireland validation cohort

Bcl-xL was evaluated within a Northern Ireland population-based cohort of stage II/III CC patient samples (n=740) using immunohistochemical methods. The Northern Ireland Cancer Registry was used to identify all patients who underwent surgery in Northern Ireland between 2004 and 2008, for a single, primary, stage II or III colon adenocarcinoma (n=1,539). This cohort was assembled into a tissue microarray, containing 3 cores from epithelial-rich tumor regions per patient. Mutational analysis was undertaken for KRAS and BRAF on n=661 (89%) of the TMA cohort using the ColoCarta panel (Agena Bioscience, Hamburg, Germany). Following sequencing, mutational status of *BRAF* and *KRAS* was available for a sub-cohort (n=661; *BRAFMT* n=92, *KRASMT* n=248, *WT/WT* n=321). We assessed *Bcl-xL* protein expression using digital pathology software, QuPath, [5] to give a numerical representation of both the extent and the intensity of staining (H-score), based on the mean expression of all cores (3 cores for each patient). All scoring was performed while blinded to the clinical details and survival endpoints. We assigned patients in each mutational genotype into high, medium or low tertile groups according to their Bcl-xL protein expression H-score.

#### Statistical analysis

Cox Proportional hazards analysis was conducted for both the transcriptional dataset and Northern Ireland cohort, prior to and after adjustment for potential confounders, to evaluate the association between *Bcl-xL* and survival in CC patients, according to *BRAF* and *KRAS* mutation status (Stata version 11.2, StataCorp, College Station, TX, USA).

Additional details are available in supplementary methods section.

## Results

### Study outline and rationale for risk stratification in BRAFMT CC

We analyzed available transcriptional data from the well-characterized dataset, GSE39582 (Supplementary Figure 1). Compared to *KRASMT* and *WT/WT*, *BRAFMT* patients were significantly more likely to be older (p<0.001), proximal (p<0.001) exhibit microsatellite instability (MSI, p<0.001) and be assigned as Consensus Molecular Subtype 1 (CMS1, p<0.001) (Table 1). Additionally, patients with *BRAFMT* tumors were significantly more likely to be female (p=0.04 and p=0.001) and receive no adjuvant chemotherapy (p=0.001 and p=0.006) compared to *KRASMT* and *WT/WT* respectively (Supplementary Table 1). Finally, *BRAFMT* patients were significantly more likely to have earlier stage disease (stage II v III) compared to *WT/WT* patients (p=0.04) (Supplementary Table 1). Using the 64 gene *BRAF* classifier identified by Popovici *et al.*[2] we performed semi-supervised hierarchical clustering of the gene expression profiles of the entire stage II/III patient cohort. We identified a subgroup accounting for 28% (n=127) of the tumor profiles using this method of clustering, which displayed an expression pattern similar to the pred-BRAFm profile (Supplementary Figure 2A). We confirmed no difference in relapse rates between the pred-BRAFm and the pred-BRAFwt populations in this cohort (Supplementary Figure 2A; HR=0.95 (95% CI 0.65-1.39)).

**Table 1:** Unadjusted and adjusted analyses of relapse-free survival. RFS analysis was performed using Cox proportional hazards method in the *BRAFMT*, *KRASMT* or *WT/WT* stratified by *Bcl-xL* expression levels. Analysis was performed both before and following adjustment. *Cut-offs for low/medium/high *Bcl-xL* gene expression based on tertile values within each BRAF/KRAS status subgroup. **Adjustments included age and sex, and were tested for TNM stage, MSI status, adjuvant chemotherapy receipt and tumor location for all models. A backwards elimination model was applied for tested confounders, until all were significant at the p<0.25 level in the model. Final adjustments included age, sex, TNM stage and MSI status (for BRAF MT and WT/WT); age, sex, TNM stage, adjuvant chemotherapy receipt and tumor location (for KRAS MT).

| **Bcl-xL** | **Non-progressors**<br>**n** | **Progressors**<br>**n** | **Unadjusted Hazard ratios**<br>**(95% confidence intervals)** | **Adjusted\*\* Hazard ratios**<br>**(95% confidence intervals)** |
| --- | --- | --- | --- | --- |
| **BRAF MT** |  |  |  |  |
| **Low** | 14 | 0 | 1.00 | 1.00 |
| **Medium** | 11 | 2 | 1.80 (Not calculable) | 3.94 (Not calculable) |
| **High** | 8 | 6 | 5.83 (Not calculable) | 9.63 (Not calculable) |
| **KRAS MT** |  |  |  |  |
| **Low** | 38 | 18 | 1.00 | 1.00 |
| **Medium** | 33 | 21 | 1.45 (0.75-2.77) | 1.47 (0.76-2.84) |
| **High** | 34 | 22 | 1.25 (0.65-2.40) | 1.32 (0.68-2.56) |
| **WT/WT** |  |  |  |  |
| **Low** | 57 | 13 | 1.00 | 1.00 |
| **Medium** | 53 | 17 | 1.39 (0.67-2.85) | 1.27 (0.61-2.64) |
| **High** | 50 | 20 | 1.54 (0.77-3.10) | 1.47 (0.72-3.01) |
| **MT: Mutant; WT/WT: BRAF and KRAS wild-type.**<br>**\*Cut-offs for low/medium/high Bcl-xL gene expression based on tertile values within each BRAF/KRAS status subgroup.**<br>**\*\*Adjustments included age and sex, and were tested for TNM stage, MSI status, adjuvant chemotherapy receipt and tumour location for all models. A backwards elimination model was applied for tested confounders until all were significant at the p<0.25 level in the model. Final adjustments included age, sex, TNM stage and MSI status (for BRAF MT and WT/WT); age, sex, TNM stage, adjuvant chemotherapy receipt and tumour location (for KRAS MT).** |  |  |  |  |

### Gene expression associated with risk of relapse in BRAFMT CC

GSEA of the discovery subset indicated increased myogenesis, epithelial-to-mesenchymal transition (EMT) and hypoxia pathways in the *BRAFMT* tumors with the highest-risk of disease relapse (Supplementary Fig 2B). Additionally, using the MCP-counter, we identified a non-significant trend for increased fibroblasts in high-risk *BRAFMT* tumors (Supplementary Fig 2C). Using differential gene expression analysis contrasting profiles from high-risk or low-risk *BRAFMT* tumors in the discovery subset (Supplementary Figure 1), we identified 83 probesets (Supplementary Table 2) corresponding to 67 annotated genes that are prognostic for relapse risk in *BRAFMT* tumors; high expression of 43 genes were associated with increased risk of relapse, and high expression of 24 genes with decreased risk of relapse (Supplementary Table 3). Increased expression of endoplasmic reticulum stress-induced transcripts such as PPP1R15A (GADD34), heat shock proteins HSPA6 and DNAJB1, and the stress-related transcription factor DDIT3 were observed in *BRAFMT* tumours with the highest-risk of disease relapse.

While the majority of the 67 genes are represented by a single probeset, *BCL2L1* (encoding *Bcl-xL*) and *NCRNA00275* (which transcribes *ZFAS1*) are both represented by 3 *different* probesets, reducing the probability of the single genes themselves being false positives, which could potentially confound the validity of genes identified by a single probeset only (Supplementary Table 2). Gene expression levels of *Bcl-xL* were increased between 1.76-1.97 fold (Figure 1A) and *ZFAS1* by 1.83 – 1.90 fold (Supplementary Table 2) in the high-risk group compared to the low risk group. Importantly, the 67 *BRAFMT* prognostic gene list is distinct from the *BRAFMT* transcriptional classifier reported by Popovici, as only one gene, (Kallikrein-Related Peptidase 10 (KLK10)), overlaps between these 2 gene lists (Supplementary Figure 2D).

### Probesets associated with risk in BRAFMT tumors represent distinct novel prognostic biology

As *BRAF* and *KRAS* are both key components of the EGFR/MAPK pathway, we performed a similar risk association analysis in *KRASMT* tumors and identified 139 probesets associated with risk-of-relapse in this genetic subgroup (Supplementary Table 4). We found no overlap between the probesets associated with risk-of-relapse in the *KRASMT* subgroup and the probesets identified in the *BRAFMT* analyses (Supplementary Figure 2E), indicating that distinct biologies determine prognosis in these two subgroups, at least in stage II/III disease.

### Bcl-xL mRNA expression is associated with poor prognosis in stage II/III BRAFMT CC

In order to confirm the genotype-specific prognostic relevance of elevated *Bcl-xL* gene expression for *BRAFMT* CC, we next generated an “Initial Consolidation” dataset (n=417, Supplementary Figure 1) by removing the filters initially applied to the discovery subset of the GSE39582 cohort, (i.e. we removed the restrictions on chemotherapy treatment and the follow-up criteria as detailed in Methods). This set of 417 patients represents an ideal cohort to assess the prognostic value of *Bcl-xL* in *KRASWT* and *WT/WT* patients that were not used to identify *Bcl-xL*, in addition to a further 17 *BRAFMT* patients that were previously excluded from the discovery data. Within *BRAFMT* tumors (n=41), the *Bcl-xL*-high group (*Bcl-xL*^high^) had a significantly higher risk-of-relapse compared to the *Bcl-xL*-low (*Bcl-xL*^low^) expression group, as determined using an unadjusted model (HR=5.83). In line with established clinical findings that MSI tumors are less likely to relapse in early stage disease[3], we performed an adjusted model (HR=9.63) accounting for potential confounding factors including age, gender, TNM stage and MSI status (confidence intervals could not be calculated due to an absence of events in *Bcl-xL*^low^; Figure 1B and Table 1). When examining untreated patients only, the prognostic value of *Bcl-xL* mRNA expression in *BRAFMT* patients was again apparent (Figure 1C); however, the prognostic value of *Bcl-xL* in the chemotherapy-treated patient subgroup could not be evaluated due to small numbers (n=8). Stratification based on the median also demonstrated the prognostic value of *Bcl-xL* gene expression (HR=5.24 (95% CI 1.3-21.2)) (Supplementary Figure 3A).

In contrast, although there was a suggestive prognostic trend, no significant associations were observed for *Bcl-xL* gene expression in either the *KRASMT* or *WT/WT* patient groups, as assessed using either an adjusted or unadjusted analysis model (Table 1 and Supplementary Figures 3B and 3C).

### ZFAS1 mRNA expression is associated with poor prognosis in stage II/III BRAFMT CC

High gene expression of *ZFAS1* was associated with a prognostic trend in *BRAFMT* tumors compared to low gene expression (Supplementary Figure 4A), although this trend failed to reach significance in either unadjusted HR=4.69 (95% CI 0.52-42.01), or adjusted HR=4.71 (95% CI 0.50-44.00) analyses (Supplementary Table 5). There was no prognostic value associated with high *ZFAS1* expression in the *KRASMT* population (adjusted HR=0.65 (95% CI 0.34-1.24)), although there was a significant association with lower relapse rates in the WT/WT population (adjusted HR=0.47 (95% CI 0.24-0.92)) (Supplementary Figure 4, Supplementary Table 5).

### Bcl-xL gene and protein expression are associated with the epithelial component of the tumor

Given the multiple cell types that constitute the tumor microenvironment (TME) in CC, we analyzed *Bcl-xL* mRNA expression levels in transcriptional data derived from micro-dissected tumor tissue (GSE35602). We observed that its expression was bimodal in the epithelial compartment of the TME, with high and low subgroups around the median, whereas stromal expression levels were generally low, with values below the median (Supplementary Figure 5A). Analysis of matched Bcl-xL transcript abundance (by Agilent microarray) and protein levels (by Reverse Phase Protein Array (RPPA)) from 102 CRC tumor samples within The Cancer Genome Atlas (TCGA)[6] indicated a significant correlation between both methodologies (p=0.001; Supplementary Figure 5B), supporting protein-based assessment as an appropriate methodology to validate our data in an independent cohort. Following optimization of an IHC protocol for *Bcl-xL* protein expression, the predominantly epithelial-derived nature of *Bcl-xL* protein expression and neoplastic-specific staining compared to the normal glands in surrounding tissue was confirmed in a series of whole-face CC sections, although there does appear to be some stromal expression, in line with our transcriptional analysis (Figure 2A).

**Figure 2:**
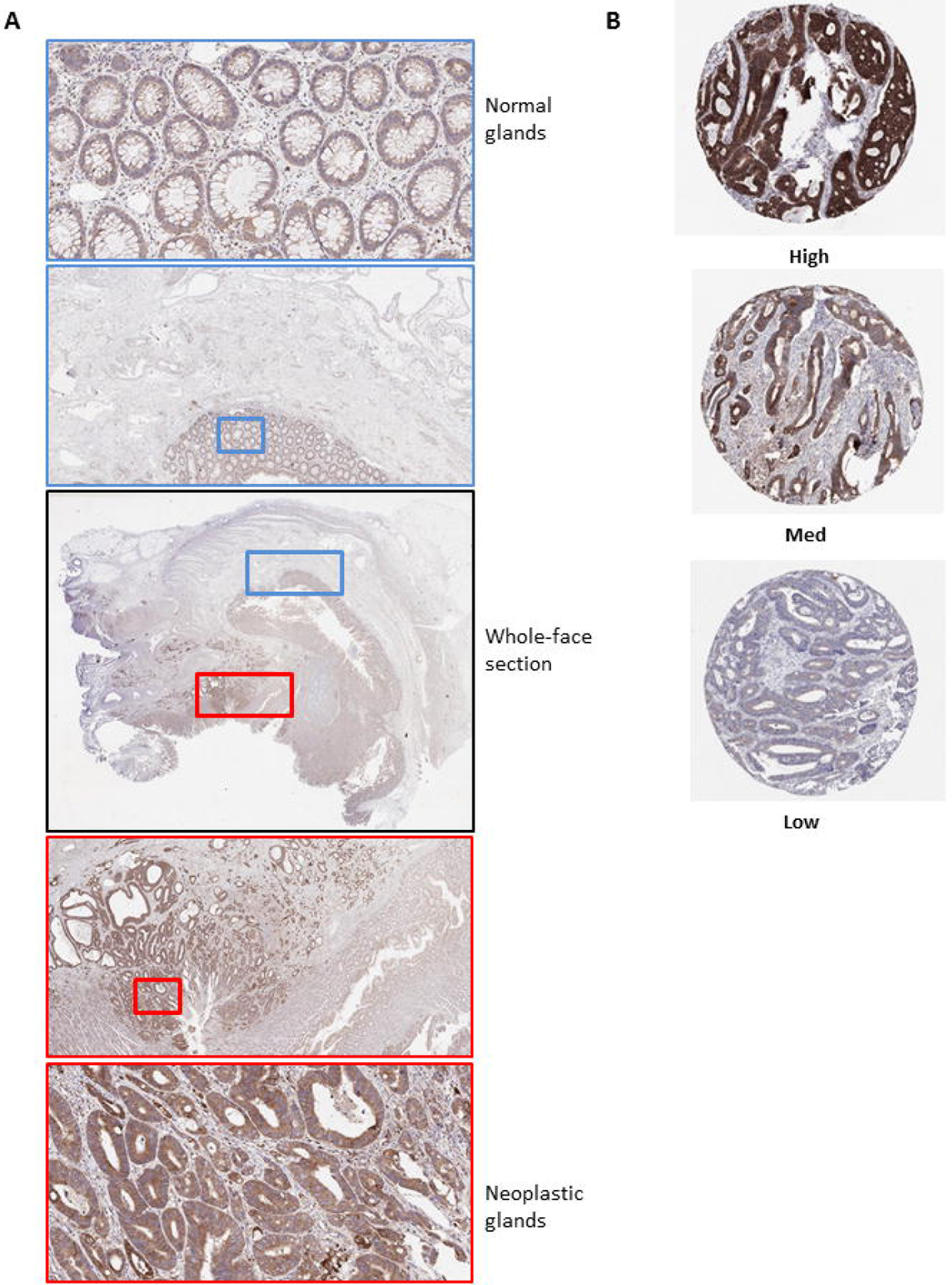
Optimization of immunohistochemistry staining protocol for Bcl-xL protein expression in CC. **A.** Whole-face CC tissue sections were used to optimize IHC protocol. A low level of protein expression was observed in the normal glands compared to surrounding stroma (Blue box) Elevated levels of expression were observed in neoplastic glands compared to both the normal glands and surrounding stroma (Red box). Some staining in the stroma is evident in both normal and cancer-associated regions. **B.** Representative images of high, medium and low Bcl-xL protein expression by IHC in an independent “Northern Ireland cohort” of stage II/III CRC (Northern Ireland cohort; n=740).

### Independent validation of Bcl-xL as a poor prognostic marker specifically in stage II/III BRAFMT CC

We then independently validated the prognostic value of *Bcl-xL* protein expression specifically in *BRAFMT* patient samples from within a Northern Ireland cohort (n=661) (Supplementary Figure 1 and described in Methods). Employing tertiles defined by protein expression (Figure 2B), we found that in *BRAFMT* disease *Bcl-xL*^high^ was associated with an increased risk of CRC disease-specific survival (DSS) when compared with *Bcl-xL*^low^, in both unadjusted (HR=3.07 (95% CI 1.24-7.60)) and using an adjusted model to account for MSI status (HR=5.50 (95% CI 1.71-17.69) (Supplementary Figure 6A and Table 2). Similar findings were evident when using OS as the endpoint (Supplementary Figure 6B).

We next conducted stratified analyses within this cohort to assess independently the prognostic value of Bcl-xL protein expression in both untreated and chemotherapy-treated *BRAFMT* patients. In untreated patients, we observed a 12-fold increased DSS risk in patients with the highest Bcl-xL protein expression (adjusted model accounting for MSI HR=12.13 (95% CI 2.49-59.13)) (Figure 3A), which was not observed in treated patients, (adjusted model accounting for MSI HR=0.96 (95% CI 0.08-11.42)) (Supplementary Figure 6C and Table 2). This significant prognostic benefit from adjuvant chemotherapy in *BRAFMT* patients was only observed in patients with the highest Bcl-xL protein expression (*P*-value for interaction =0.006), whereas patients with low Bcl-xL protein expression derived no benefit from the addition of chemotherapy (Figures 3B and C, Table 2 and Supplementary Figure 6C). Similar results were evident when using OS as the endpoint (Supplementary Figure 6D, E and F). Importantly, in agreement with our initial consolidation cohort, we were again able to confirm that the prognostic value of Bcl-xL protein expression was not observed in *KRASMT* (HR=1.00 (95% CI 0.57-1.77) and *WT/WT* (HR=1.18 (95% CI 0.67-2.09)) patient samples (Table 4).

**Figure 3:**
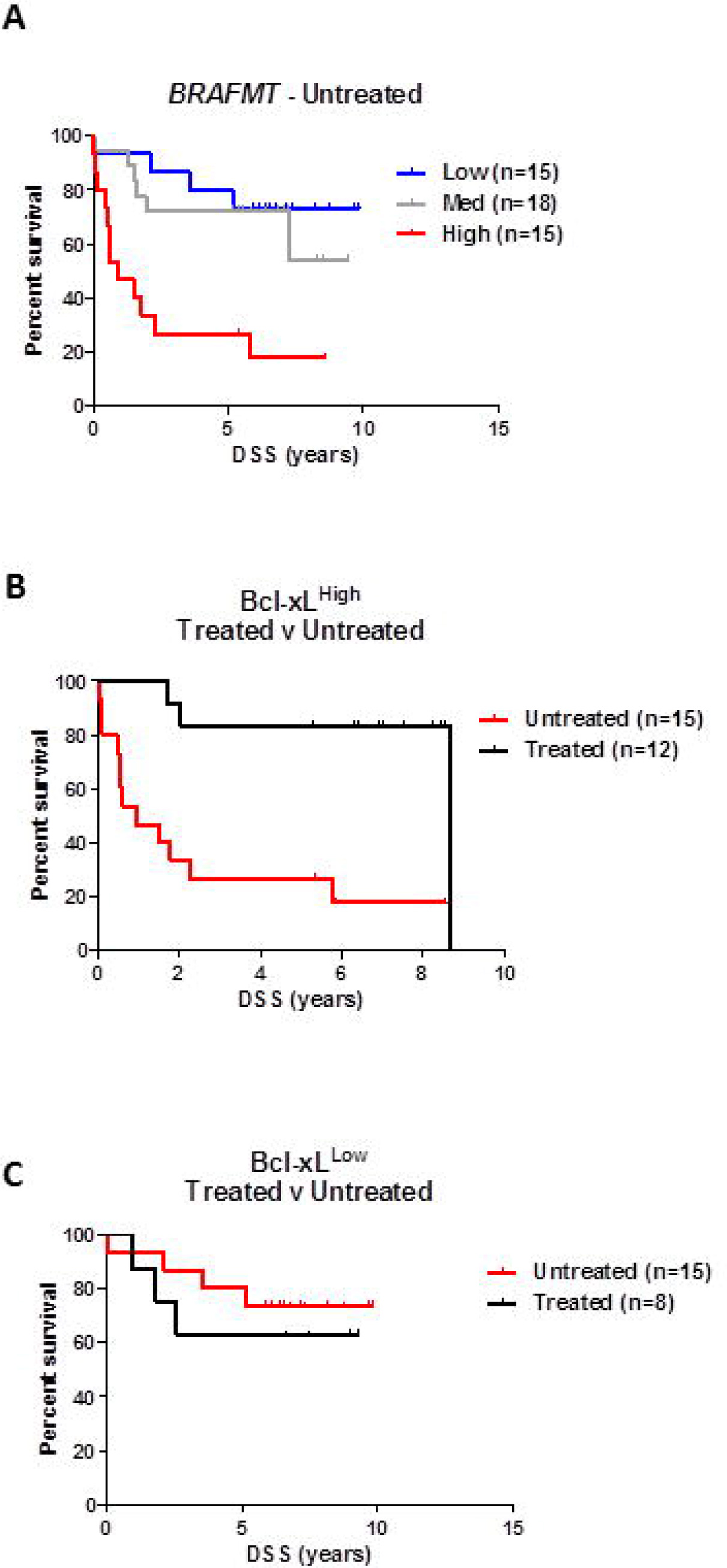
Independent validation of the prognostic value of Bcl-xL protein expression in *BRAFMT* CC. **A.** Colorectal cancer disease-specific survival (DSS) curve using Kaplan-Meier estimation in the “Northern Ireland cohort” comparing tertiles stratification of *Bcl-xL* protein expression (by IHC H-score) in untreated *BRAFMT* stage II/III CC patients. **B.** DSS of patients in the highest tertile of *Bcl-xL* protein expression stratified according to adjuvant chemotherapy treatment received. **C.** DSS of patients in the lowest tertile of *Bcl-xL* protein expression stratified according to adjuvant chemotherapy treatment received. Unadjusted and adjusted HR statistics are detailed in Table 4.

**Table 2:** Analyses of disease-specific survival in the independent IHC validation cohort. **(Top)** DSS analysis was performed using Cox proportional hazards method in the *BRAFMT*, *KRASMT* or *WT/WT* stratified by Bcl-xL IHC (H-score) protein expression levels. Analysis was performed both before and following adjustment. *Cut-offs for low/medium/high *Bcl-xL* gene expression based on tertile values within each BRAF/KRAS status subgroup. **Adjustments included age, sex, TNM stage, MSI status, adjuvant chemotherapy receipt, ECOG status, family history of colorectal cancer, year of diagnosis and extramural venous invasion for all models. **(Bottom)** Further adjusted analysis to identify treatment interaction effect of the Bcl-xL-high tertile group of BRAFMT tumors stratified by treatment received.

| Bcl-xL | Alive<br>n | CRC Death<br>(DSS) N | Unadjusted Hazard ratios<br>(95% confidence intervals) | Adjusted** Hazard ratios<br>(95% confidence intervals) |  |  |
| --- | --- | --- | --- | --- | --- | --- |
| BRAF MT |  |  |  |  |  |  |
| Low (<56.1) | 16 | 7 | 1.00 | 1.00 |  |  |
| Medium (56.1-<91.1) | 17 | 10 | 1.54 (0.58-4.09) | 1.97 (0.60-6.44) |  |  |
| High (≥91.1) | 12 | 15 | 3.07 (1.24-7.60) | 5.50 (1.71-17.69) |  |  |
| KRAS MT |  |  |  |  |  |  |
| Low (<53.5) | 44 | 28 | 1.00 | 1.00 |  |  |
| Medium (53.5-<92.7) | 41 | 24 | 0.98 (0.57-1.69) | 0.93 (0.52-1.66) |  |  |
| High (≥92.7) | 38 | 33 | 1.37 (0.82-2.27) | 1.00 (0.57-1.77) |  |  |
| WT/WT |  |  |  |  |  |  |
| Low (<66.1) | 57 | 28 | 1.00 | 1.00 |  |  |
| Medium (66.1-<105.8) | 58 | 29 | 0.99 (0.59-1.68) | 1.05 (0.60-1.84) |  |  |
| High (≥105.8) | 59 | 31 | 1.07 (0.64-1.78) | 1.18 (0.67-2.09) |  |  |
| MT: Mutant; WT/WT: BRAF and KRAS wild-type.<br>*Cut-offs for low/medium/high Bcl-xl gene expression based on tertile values within each BRAF/KRAS status subgroup.<br>**Adjustments included age, sex, TNM stage, MSI status, adjuvant chemotherapy receipt, ECOG status, family history of colorectal cancer, year of diagnosis and extramural venous invasion for all models. |  |  |  |  |  |  |
| Bcl-xL | No Chemotherapy receipt |  |  | Chemotherapy receipt |  |  |
|  | Alive<br>n | CRC Death<br>(DSS)<br>N | Adjusted Hazard ratios<br>(95% confidence<br>intervals) | Alive<br>n | CRC Death<br>n | Adjusted Hazard ratios<br>(95% confidence<br>intervals) |
| BRAF MT |  |  |  |  |  |  |
| Low (<56.1) | 11 | 4 | 1.00 | 5 | 3 | 1.00 |
| Medium (56.1-<91.1) | 12 | 6 | 1.99 (0.38-10.29) | 5 | 4 | 2.18 (0.23-20.89) |
| High (≥91.1) | 3 | 12 | 12.13 (2.49-59.13) | 9 | 3 | 0.96 (0.08-11.42) |
| P for interaction |  |  | 0.006 |  |  |  |
| MT: Mutant.<br>Cut-offs for low/medium/high Bcl-xl gene expression based on tertile values within each BRAF MT subgroup.<br>Adjustments included age, sex, TNM stage, MSI status, adjuvant chemotherapy receipt, ECOG status, family history of colorectal cancer, year of diagnosis and extramural venous invasion. |  |  |  |  |  |  |

## Discussion

In this study, we set out to identify factors influencing patient prognosis specifically in tumors harboring an oncogenic *BRAF* mutation. Stratification of a discovery prognostic cohort based on risk-of-relapse identified the *Bcl-2* family member, *Bcl-xL*, as being upregulated at the transcriptional level in *BRAF*MT tumors from patients who went on to relapse following surgery, compared to those *BRAFMT* patients who experienced no disease recurrence. We validated the prognostic value of *Bcl-xL* specifically in *BRAFMT* tumors in both a consolidation transcriptional cohort and in an independent population-based stage II/III cohort. Importantly, in each validation series, we also confirmed the *BRAFMT*-specific nature of this association, as the expression of *Bcl-xL* was not associated with increased risk of disease relapse or death in either *KRASMT* or *WT/WT* tumors. Interestingly, we observed that although *BRAFMT* tumors with high *Bcl-xL* expression have a poor prognosis, this subgroup also appears to benefit the most from standard adjuvant chemotherapy.

The prognostic value of stratifying CC patients based on *BRAF* mutational status has been well reported, particularly in stage IV metastatic tumors, where patients with *BRAFMT* tumors have poor survival rates. A previous study identified a transcriptional signature that could stratify stage II and III CRC tumor profiles into subgroups based on their similarity to *BRAFMT* tumors (pred-BRAFm).[2] The authors demonstrated the utility of either *BRAF* mutational status or the pred-BRAFm classifier in identifying patients with shorter survival, although no difference was observed in the initial disease-specific relapse rates between the identified subgroups. This important result suggests that the prognostic power associated with the pred-BRAFm signature, or indeed the presence of the *BRAF* mutation itself, is due to shorter survival time because of aggressive disease <u>*after*</u> relapse in stage IV disease; however, initially, *BRAFMT* stage II and III patients are not at a higher risk of their early-stage disease progressing to metastatic disease. This subtle but crucial point underpins our rationale for performing a stratified analysis to identify factors determining risk-of-relapse specifically <u>*within*</u> the *BRAFTMT* genotype. The data presented here identifies for the first time a novel role for *Bcl-xL* expression in influencing disease relapse, providing a new, important and clinically relevant understanding of the biology underpinning aggressive *BRAFMT* stage II/III disease. Interestingly, we found almost no overlap between the genes associated with relapse in *BRAFMT* and *KRASMT* tumors, suggesting that although there is constitutive activation of the MAP kinase pathway in both these subgroups, there is clearly distinct prognostic tumor biology associated with these different genotypes.

A recent study using RPPA methodology reported that a mathematical model of Bcl-2 family protein interactions (including Bcl-xL) termed DR_MOMP was prognostic in chemotherapy-treated stage III CRC.[7] Moreover, this study found that Bcl-2 family signaling was particularly important in Consensus Molecular Subtypes (CMS) 1 and 3. As the CMS1 subgroup is enriched for *BRAFMT* disease, this report appears to be in agreement with our current study. However, the individual contribution of Bcl-xL expression to prognosis in CMS1/*BRAFMT* CRC was not reported.

The reason for the significant benefit from standard chemotherapy of Bcl-xL high *BRAFMT* CRC is unclear. High Bcl-xL expressing tumors may be “primed” to undergo apoptosis in response to chemotherapy, due to co-expression of pro-apoptotic Bcl-2 family members.[8, 9]. The findings presented both here and by others suggest that *BRAFMT* driven CRC is more aggressive than *BRAFWT* disease, but only when the disease has disseminated from the primary site. Interestingly, we observed specific changes in the ER-stress machinery in *BRAFMT* tumors with the highest-risk of disease relapse, with activation and upregulation of factors including GADD34, heat shock proteins, and stress-related transcription (DDIT3) in our analysis. Additionally, using GSEA, we identify increased hypoxia and EMT signaling in high-risk tumors, again indicating an association with ER stress-activation. Each of these factors have been demonstrated to activate the unfolded protein response (UPR), which in turn has been correlated with a higher risk of metastatic recurrence in breast cancer.[10, 11] In agreement with our findings, upregulation of UPR signaling in disseminated tumor cells from breast cancer, lung cancer and prostate cancer enables both the formation and long-term persistence of metastatic lesions.[10, 11] In addition to activation of the UPR machinery, high Bcl-xL expression may promote survival of invasive tumor cells during the metastatic process; for example Bcl-xL has been reported to be a suppressor of anoikis,[9, 12] which would explain its association with increased risk-of-relapse in the *BRAFMT* subgroup.

This study has several strengths. We have identified Bcl-xL as a novel predictor of response within a poor prognostic group of CC patient samples, using a robust approach that included validation in an independent cohort using a clinically relevant methodology. The population-based nature of our validation cohort also means that the results should be generalizable to all stage II/III CRC patients. Additionally, given that IHC and mutational tests for *BRAF* and *KRAS* are routinely utilized in the diagnostic work-up of CC patients, the methods we have used here could easily be employed within routine pathology reporting practice. However, we do acknowledge that further work is required to identify an optimal cut-off level of *Bcl-xL* expression that would allow a more robust classification of low and high expressers for prospective patient stratification.

In conclusion, we have identified and independently validated the prognostic value of Bcl-xL mRNA and protein expression specifically within *BRAFMT* CC, which should help inform selection of treatment options for high-risk *BRAFMT* stage II/III patients in the adjuvant disease setting. This approach could prevent the initial relapses, which ultimately contribute to the poor outcomes of patients with this genotype. Data presented here provide compelling evidence that, in addition to *BRAF* mutational analysis, assessment of Bcl-xL protein expression using routine diagnostic IHC methods can identify both poor prognostic *BRAFMT* stage II/III CC patients who will benefit from adjuvant therapy and an otherwise good prognostic subgroup of *BRAFMT* patients who derive no significant advantage from the addition of adjuvant chemotherapy.

## Acknowledgements

Dr Philip Dunne has received funding from Andrew Entwistle, in the form of a travel grant, enabling a visiting fellowship at the University of Torino and the completion of this work. The samples used in this research were received from the Northern Ireland Biobank which is funded by HSC Research and Development Division of the Public Health Agency in Northern Ireland and Cancer Research UK through the Belfast CR- UK Centre and the Northern Ireland Experimental Cancer Medicine Centre; additional support was received from the Friends of the Cancer Centre. The Northern Ireland Molecular Pathology Laboratory which is responsible for creating resources for the NIB has received funding from Cancer Research UK, the Friends of the Cancer Centre and the Sean Crummey Foundation. The Northern Ireland Cancer Registry is funded by the Public Health Agency, Northern Ireland. We also thank Ken Arthur (Belfast) for his expertise in TMA construction of the cohort, Enzo Medico (Torino) for his advice and expertise, and all individuals who were involved in the creation and study design of the Northern Ireland cohort.

## Specific author contributions

**PDD:** Design of the study, data acquisition and analysis, interpretation of results, drafting the work and study supervision; **HGC:** Data acquisition and analysis, interpretation of results and revision of article; **PB**, **MA:** Data acquisition and analysis; **RTG:** Data acquisition, interpretation of results, collection of samples; **SMcQ, VB, MBL, JAJ:** Collection of samples; **AMBMcC, AG, DK:** Data analysis; **CH, FDiN, ST:** Interpretation of results; **DMcA, PGJ, DBL:** Study supervision, interpretation of results; **ML:** Design of the study, interpretation of results, revision of article, study supervision and funding. All authors approved the final version of the submitted work.

